# COVID-19 induces persistent transcriptional changes in adipose tissue that are not associated with Long COVID

**DOI:** 10.1101/2025.05.23.655815

**Authors:** Soneida DeLine-Caballero, Kalani Ratnasiri, Heping Chen, Seynt Jiro Sahagun, Uma M. Mangalanathan, Trisha R. Barnard, Nicole Turk, Jasmine Yang, Natesh Saini, Ekrem M. Ayhan, Hasiyet Memetimin, Brian S. Finlin, Zachary Leicht, Phillip A. Kern, Ivette Emery, Brooke M. Leeman, Gregory E. Edelstein, Samantha Costa, Alina Choi, Jonathan Z. Li, Clifford Rosen, Tracey McLaughlin, Catherine A. Blish

## Abstract

Long COVID is a heterogeneous condition characterized by a wide range of symptoms that persist for 90 days or more following SARS-CoV-2 infection. Now more than five years out from the onset of the SARS-CoV-2 pandemic, the mechanisms driving Long COVID are just beginning to be elucidated. Adipose tissue has been proposed as a potential reservoir for viral persistence and tissue dysfunction contributing to symptomology seen in Long COVID. To test this hypothesis, we analyzed subcutaneous adipose tissue (SAT) from two cohorts: participants with subacute COVID-19 (28–89 days post-infection) compared to pre-pandemic controls, and participants with Long COVID compared to those with those classified as “indeterminate” based on the RECOVER-Adult Long COVID Research Index (12-47 months post-infection). We found no evidence of persistent SARS-CoV-2 RNA in adipose tissue in any participant. SAT from participants with subacute COVID-19 displayed significant transcriptional remodeling, including depleted immune activation pathways and upregulated Hox genes and integrin interactions, suggesting resident immune cell exhaustion and perturbations in tissue function. However, no consistent changes in gene expression were observed between Long COVID samples and samples from indeterminant participants. Thus, SAT may contribute to inflammatory dysregulation following COVID-19, but does not appear to play a clear role in Long COVID pathophysiology. Further research is needed to clarify the role of adipose tissue in COVID-19 recovery.

## 2 Introduction

Long COVID (LC) is a complex heterogeneous disease characterized by a wide variety of symptoms that persist 90 days or more after a SARS-CoV-2 infection. LC can manifest as neurological, cardiovascular, gastrointestinal, psychiatric, or pulmonary symptoms; however, the disease has also been attributed to more systemic features such as brain fog and chronic fatigue (1). As of April 6th, 2025 the World Health Organization has reported 777,704,325 cases of acute coronavirus disease 2019 (COVID-19), and thousands of new COVID-19 cases continue to be reported each week (2). The United States Centers for Disease Control estimates that approximately 6.9% of all people who become infected with SARS-CoV-2, regardless of their symptom severity, will experience disease progression into LC (3).

While the exact mechanisms driving LC remain unknown, potential factors driving symptomatology include persistent SARS-CoV-2 replication, inflammation, and/or overall dysfunction in an unknown tissue reservoir(1,4–10). Early in the COVID-19 pandemic, numerous findings pointed to adipose tissue’s potential as a reservoir for disease. Adipose tissue shapes systemic inflammation and metabolic states through the release of adipose specific chemokines, adipokines, and metabolites (11). Therefore, disturbances in adipose tissue function, as seen in obesity and other metabolic comorbidities, can have systemic consequences throughout the body (11). Indeed, obesity has emerged as one of the strongest independent risk factors for developing severe acute COVID-19 and persistent symptoms documented in LC (12–14). Dysfunctional, pro-inflammatory obese adipose tissue plays a critical role in the development of systemic insulin resistance which leads to Type 2 Diabetes (15). Several prospective studies have followed cohorts 3-9 months after disease onset and found that there is an increased risk of a Type 2 diabetes diagnosis in pediatric and adult participants which could suggest that SARS-CoV-2 remodels the function of adipose tissue (16–19). In addition, hyperglycemia and low adiponectin levels were noted in individuals infected with SARS-CoV-2 (20).

Several studies suggested that infection of adipose tissue could contribute to this metabolic dysfunction. SARS-CoV-2 infected adipose tissue in a hamster model and infected human adipocytes *in vitro (21,22)*. Our group found that both adipocytes and resident macrophages isolated from subcutaneous and visceral adipose tissue were infected with SARS-CoV-2 *in vitro*, inducing pro-inflammatory cytokines such as IP-10 and MCP-2 (12). Autopsy studies have found markers of viral persistence in adipose tissue collected from hospitalized patients whose death was attributed to severe acute COVID-19 in the early years of the pandemic (12,23). Though these prior investigations have examined viral persistence in adipose tissue in the context of fatal cases of acute COVID-19, little is known about adipose tissue remodeling and viral persistence in living individuals.

We hypothesized that persistent infection of adipose tissue might contribute to LC. To investigate this, we profiled 27 participants across two separate cohorts of living COVID-19 participants: in the subacute phase (28-90 days post positive COVID-19 test), and in the LC phase (90+ days post positive COVID-19 test). We performed quantitative RT-PCR for SARS-CoV-2 N gene and bulk RNA-sequencing to determine whether SARS-CoV-2 was present in adipose tissue, whether COVID-19 infection was associated with changes in the inflammatory profile of adipose tissue, and whether changes in adipose tissue were associated with LC symptoms.

## 3 Materials and methods

### 3.1 Study Design

Cohort 1 consisted of 9 healthy adults without diabetes undergoing adipose tissue biopsy who consented for studies of adipose tissue inflammation/infection (NCT# 06833217), or who consented to participate in the RECOVER study at Stanford University (Table 1, Supplementary Table 1). COVID-19 history was obtained, and biopsies were deferred to at least 30 days after infection for those with a positive SARS-CoV-2 antigen test. Eligibility included age 30-79 yrs, BMI 25-35 kg/m^2^, stable body weight for four weeks, and absence of diabetes based on fasting glucose < 126 mg/dL. Exclusion criteria included major organ disease, chronic inflammatory conditions, malignancy, acute infection or trauma, history of bariatric surgery or liposuction, use of medications known to alter body weight, adipose tissue lipolysis, or blood glucose. The study was approved by the Stanford IRB and all participants gave written informed consent. Five participants had subacute with subcutaneous adipose tissue (SAT) samples collected 28 - 44 days after a positive COVID-19 antigen test between August and November 2023. Four non-infected participants who underwent fat biopsy pre-pandemic (March 2019 through Feb 14, 2020) were selected as controls.

**Table 1:** Summary of Clinical Data for Cohort 1 (Subacute COVID & Pre-Pandemic Controls Samples) and Cohort 2 (Long COVID & indeterminate Status Controls.) ^*^Ct Value 37 or higher is considered below the limit of detection. **0= no symptoms to 20= Most Severe (Median).

|  | Pre-Pandemic Donors (N=4) | Subacute COVID-19 Donors (N=5) | Indeterminate COVID-19 Donors (N=10) | Long COVID-19 Donors (N=8) |
| --- | --- | --- | --- | --- |
| **Age Range in Years (Median)** | 40-54 (48.5) | 36-79 (65) | 41 to 71 (60) | 41 to 75 (56) |
| **Sex** |  |  |  |  |
| **Female (%)** | 3 (75%) | 3 (60%) | 9 (90%) | 6 (75%) |
| **Male (%)** | 1 (25%) | 2 (40%) | 1 (10%) | 2 (25%) |
| **BMI Range (Median)** | 26.2 to 36.10 (32.1) | 28.9 to 35.30 (29.5) | 19.6 to 30.70 (29.9) | 24.40 to 33.70 (30) |
| **Months After Positive COVID-19 Test (Median)** | N/A | .93-1.43 (1.03) | 13.2-19.32 (22.32) | 20.52-44.07 (37.98) |
| **\*RT-PCR for SARS-CoV-2 N Gene** | All Samples Below Limit of Detection | All Samples Below Limit of Detection | All Samples Below Limit of Detection | All Samples Below Limit of Detection |
| **\*\*RECOVER-Adult Long COVID Research Index Symptom Severity Score Range** | N/A | N/A | 0 to 11 (3.5) | 12 to 18 (16) |

Cohort 2 consisted of 18 adult participants, aged 36-75 years, who were enrolled from the Researching COVID to Enhance Recovery (RECOVER) cohort for the PROMIS study (Pathobiology in RECOVER of Metabolic and Immune Status) (Table 1, Supplementary Table 1). Individuals at Maine and Kentucky sites consented for adipose tissue biopsies under a single IRB from MaineHealth PROMIS and RECOVER. Subcutaneous Adipose Tissue (SAT) samples were collected between 1 year and 3.7 years post positive COVID-19 antigen test. On the same day as the biopsy, each participant’s Long COVID symptom severity was assessed using the RECOVER-Adult Long COVID Research Index (17). Eight participants with symptom scores greater than 11 were classified as LC and eleven participants had scores at or below this threshold and were classified as “indeterminate status” (Table 1, Supplemental Table 1). Clinical data related to age, sex, BMI were also collected but not controlled across the cohort. Cohort size was not controlled and was based on sample availability.

### 3.2 Sample Collection

For Cohort 1, subcutaneous adipose tissue (SAT) was obtained under sterile conditions with administration of local anesthesia in the periumbilical region. A 14G needle and 30ml syringe was inserted and suction applied to withdraw 2g of adipose tissue as described (24). The tissue biopsy was cleaned of excess blood by PBS wash, and immediately flash frozen in liquid nitrogen and then stored at -80°C. After thawing, the tissue was lysed in TRIzol LS (Invitrogen, USA, Cat #15596018), with isolation of total RNA using chloroform extraction and isopropanol precipitation. The RNA pellet was washed with ethanol, dried, and resuspended in DEPC-treated/Nuclease Free water and quantified using Invitrogen Qubit 4 Fluorometer and Qubit RNA High Sensitivity Assay kit Thermo Fisher Scientific (Cat# Q32855).

Samples from cohort 2 were collected from abdominal SAT by full biopsy with 2 grams of tissue partitioned into three samples, one which was immediately flash frozen in -70ºC. Samples from Maine were shipped frozen to Kentucky for RNA extraction. Kentucky samples were immediately frozen after tissue biopsy. Approximately 240 mg of SAT was placed in a 2 mL tube with 700 mL Qiazol (#79306, Qiagen) and two 0.25 g stainless steel balls, and homogenized for 1 min at 1000 rpm in a SPEX sample prep 2010 Geno Grinder according to the manufacturer’s instructions. After homogenization, the volume was adjusted to 1 ml with Qiazol, 200 mL chloroform was added, the tubes were vortexed, incubated at room temperature for 5 min, and centrifuged at 12000 rpm for 15 min at 4ºC. RNA was the extracted from the aqueous phase using a RNeasy kit (#74106, Qiagen) according to the manufacturer’s instructions.

#### 3.3 Processing and sequencing of SAT samples

At least 500 ng of RNA from each SAT sample was used for cDNA library preparation and bulk RNA sequencing, which was completed by GeneWiz/Azenta. RNA samples were diluted in nuclease-free water and tested to confirm a RNA Integrity Number (RIN) value of 6 or higher using the Agilent RNA ScreenTape assay and TapeStation system. NEBNext Ultra II RNA Library Prep Kit was used for cDNA library preparation. GeneWiz/Azenta completed sequencing with PolyA selection and generated approximately 20 million paired-end reads per sample using their Illumina NovaSeq 6000 platform (Illumina, San Diego, CA, USA). A quality report and FASTQ files were provided for further analysis.

### 3.4 Immunohistochemistry

We performed immunohistochemistry staining on a subset of SAT samples from cohort 2 (Supplemental Table 2). A portion of adipose tissue was fixed in formalin, embedded in paraffin, and sectioned, and 5 mm sections were co-stained with CD86 and CD68 antibodies or CD206 and CD68 antibodies as described (25).

### 3.5 SARS-CoV-2 N gene quantification by quantitative reverse-transcription polymerase chain reaction (qRT PCR)

For cohort 1, 5 ng of total RNA was used for 1-step RT-PCR using the SuperScript™ III Platinum™ One-Step qRT-PCR Kit with ROX (Thermo Fisher Scientific Catalog: **Catalog number** 11745100). Genomic N gene quantification was done with the use of CDC qualified primers and probes amplifying the N1 region, n2019-nCoV (Stanford University Protein and Nucleic Acid (PAN) Facility) (26). PCR reactions were prepared following manufacturer’s instructions in a total volume of 25uL per reaction performed in triplicate. Ct cutoff for negative samples was above 37, as indicated by our negative control samples. The samples were analyzed on a QuantStudio 3 real-time PCR systems (Applied Biosystems) using the following parameters: (stage 1) 10 minutes at 50ºC for reverse transcription, followed by (stage 2) 3 minutes at 95ºC for initial denaturation and (stage 3) 40 cycles of 10 seconds at 95ºC, 15 seconds at 56ºC, and 5 seconds at 72ºC.

For cohort 2, the quantitative viral load reaction contained 5 µL extracted RNA, 1x TaqPath™ 1-Step RT-qPCR Master Mix, CG (Thermo Fisher Scientific catalogue number: A15300), and primers and probes amplifying the N1 region, n2019-nCoV (Integrated DNA Technologies catalogue number: 10006713). Viral load copy numbers were quantified using qPCR RNA standards in 16-fold dilutions to generate a standard curve. The assay was run in triplicate for each sample, and two non-template control wells were included as negative controls. Positive control samples with 10,000 copies/mL were generated by extracting RNA from 50 µL of the AccuPlex™ SARS-CoV-2 Reference Material Kit (Seracare catalogue number: 0505-0168). Negative controls were made using 50 µL 10X PBS Buffer pH 7.4 (Corning catalogue number: MT21040CV). One positive and one negative control were extracted and tested alongside the participant samples and were quantified using the same standard curves. The Ct cutoff for negative samples was above 38, as indicated by our negative control sample. The samples were analyzed on QuantStudio 3 real-time PCR systems (Applied Biosystems) using the following parameters: (stage 1) 2 minutes at 25ºC, (stage 2) 15 minutes at 50ºC for reverse transcription, followed by (stage 3) 2 minutes at 95ºC for initial denaturation, and (stage 4) 45 cycles of 3 seconds at 95ºC and 30 seconds at 55ºC.

### 3.6 Processing and analysis of bulk RNA-sequencing data

Raw fastq read files went through the following processing steps:

1. FASTQC was used for assessing overall sample quality (27).
2. TrimGalore was used to trim low quality reads using the following parameters “--stringency 1 --gzip --length 20 --fastqc --paired” (28).
3. Salmon was used for alignment to the human reference genome (GRCh38) using the default paired parameters(29).

Tximport was used to import salmon alignment outputs, and DESeq2 was used for further processing (30,31). Variance stabilizing transformation normalization was performed(31). DESeq2 was used for Principal Component Analysis (PCA). For cohort 1, DESeq2 was used for differential gene expression analysis using the “apeglm” method for effect size estimation(32). For cohort 2, MetaIntegrator was used for differential gene expression to take into account possible batch effects due to the two study sites(33). We utilized over-enrichment analysis to perform pathway analysis using the Blood Transcriptional Modules (BTMs)(34). We utilized published gene score methods to generate a per-sample score where we calculated the geometric mean of all genes in the specific BTM being analyzed, and scaling was performed across scores across all samples(35).

### 3.7 Processing and analysis of publicly available single cell RNA-sequencing data

Seurat v5.0.2 was used for all single-cell analysis (36). We utilized our previously processed scRNA-seq Seurat object (12), with data available under GSE208034. We subsetted on data generated from uninfected SAT samples. We performed PCA on the first 50 components, utilized Harmony for batch correction and performed UMAP on the first 50 components. We used original cell types and grouped them at a more granular level. We utilized published gene score methods to generate a per-cell score where we calculated the geometric mean of all genes in the BTM being analyzed, and scaling was performed across the scores across all cells(35).

### 3.8 Figure generation

Figures were generated in R using the ggplot2 package(37). Statistical analyses were performed as described in figure and table legends.

## 4 Results

### 4.1 Description of Cohorts

To investigate changes in the adipose tissue of living individuals with prior history of COVID-19, we enrolled two cohorts (Table 1). To understand how subacute COVID-19 impacts adipose tissue, we collected SAT from a cohort of individuals who were 0.93-1.43 months after diagnosis of a SARS-CoV-2 infection, along with SAT samples from pre-pandemic control individuals (cohort 1). To investigate whether adipose tissue is associated with LC, we enrolled cohort 2, where we collected SAT samples from individuals who were 13-44 months out from COVID-19 and were classified as either being LC or “indeterminate” status based on their symptom severity score.

### 4.2 No Evidence of SARS-CoV-2 persistence in SAT samples from subacute or LC participants

We first examined whether there was evidence for persistence of SARS-CoV-2 in either cohort by performing qRT-PCR for the SARS-CoV-2 N gene. In the subacute COVID-19 cohort all N gene Ct values measured above the limit of detection of 37.4, indicating no detection of viral RNA (Table 1). In the LC cohort, we also did not detect viral RNA (Table 1). Therefore, we found no evidence of SAT harboring SARS-CoV-2 in either subacute COVID-19 or LC cases, which is distinct from prior reports identifying viral RNA in those deceased from acute COVID-19 early in the pandemic.

### 4.3 Subacute COVID-19 Depletes Pathways Related to Immune Activity and Upregulates Pathways Related to the Hox (Homeobox) Gene Family and Integrin Cell Surface Interactions

Though we found no evidence of persistent infection, we hypothesized that COVID-19 might lead to persistent changes in the inflammatory environment in adipose tissue. We therefore performed bulk RNA-sequencing on SAT samples from cohort 1, comparing samples collected during subacute COVID-19 to those collected pre-pandemic. Samples from subacute COVID-19 differed in their expression profile from pre-pandemic controls by principal component analysis (PCA), with the first principal component largely distinguishing these populations (Figure 1A). We next explored the genes driving these differences, and identified 6,376 unique differentially expressed genes (DEGs) between subacute COVID-19 and control samples (Fig. 1B). 833 DEGS were significantly upregulated in subacute COVID-19 samples, while 5,543 DEGs were significantly downregulated. To understand the immunological context of these DEGs, we performed gene set enrichment analysis using the Blood Transcriptome Modules (BTM)(38), which mapped differentially expressed genes in our cohort to modules of co-expressed genes associated with known biological functions (Figure 2C). Overrepresentation analysis on both the upregulated DEGs and downregulated DEGs separately revealed that several BTMs related to immune activation were depleted in subacute COVID-19 participants, while all Hox BTMs were significantly upregulated as well as a module related to integrin surface interactions. Thus, prior COVID-19 infection was associated with significant changes in the transcriptional landscape of adipose tissue, even without detecting persistent viral RNA.

**Figure 1.**
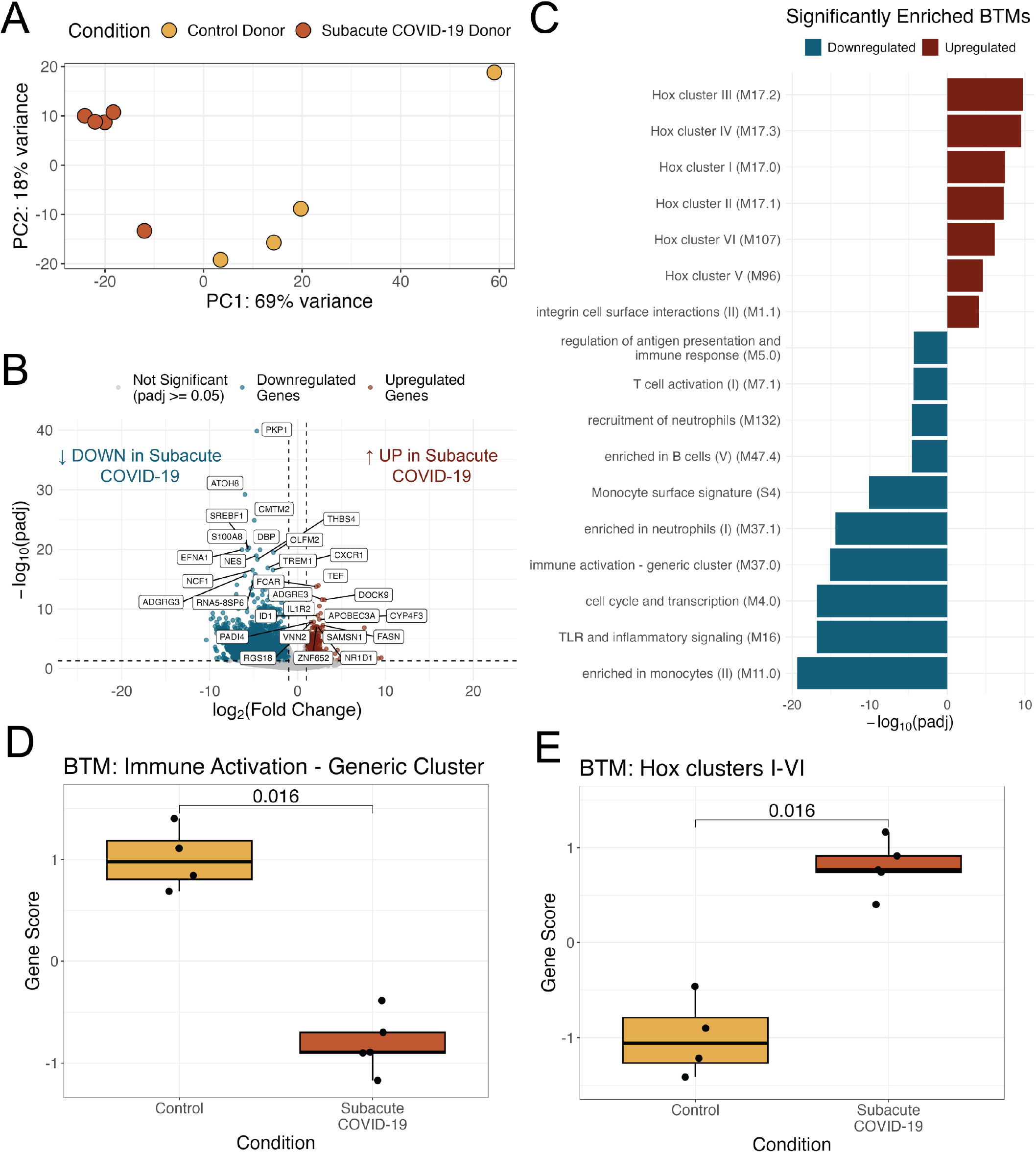
Transcriptional changes in SAT from participants with subacute COVID-19. **(A)** Dimensionality reduction by principal components (PC) analysis on all genes for participant samples in cohort 1 (subacute COVID-19 and pre-pandemic participant samples). Each data point corresponds to the SAT sample of a participant. PC1, principal component 1; PC2, principal component 2; percentage expresses contribution to the overall data variability **(B)** Volcano plot visualization of the differentially expressed genes between subacute COVID-19 participant and pre-pandemic control samples. Horizontal dashed line represents padj = 0.05 and vertical dashed lines represent an absolute log fold change of 1. **(C)** Significantly enriched (padj < 0.0001) Blood Transcriptional Modules (BTM) within the upregulated and downregulated differentially expressed genes. **(D)** Box plot of Immune activation gene scores for subacute COVID-19 samples (n=5) and pre-pandemic control samples, (n=4); p-values generated with Wilcoxon rank sum test. **(E)** Box plot Hox Cluster III BTM Gene Scores for subacute COVID-19 participants (n=5) and pre-pandemic control participants (n=5); p-values generated by Wilcoxon rank sum test. For boxplot visualizations, the top box line indicates the 75th quartile gene score; middle box line indicates the mean gene score, and the bottom box line represents the 25th quartile gene score

**Figure 2.**
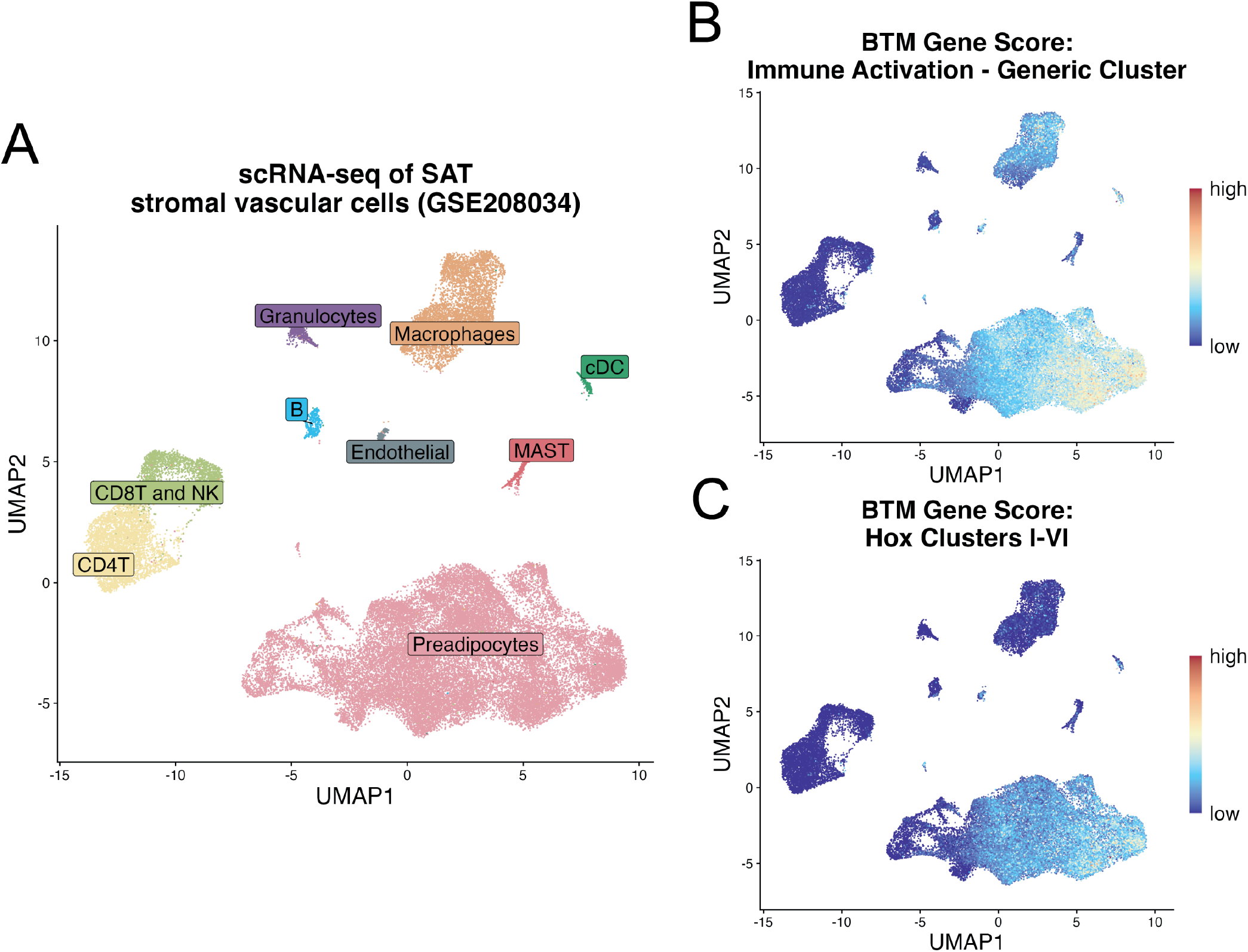
Expression of Immune Activation and Hox Cluster BTMs Are Strongly Associated with Resident Tissue Macrophages and Preadipocytes in SAT. **(A)** Uniform Manifold Approximation and Projection (UMAP) of Single Cell RNA (scRNA) dataset from Stromal Vascular Fraction of human SAT (GSE208034) colored by cell type (annotated using Seurat) **(B-C)** UMAP from **A**, colored by gene score generated on the expression of genes within the Blood Transcription Modules (BTMs) of **(B)** immune activation and **(C)** Hox clusters I-VI.

To visualize the participant-level transcriptional changes, we generated BTM gene scores, based on the geometric mean of the expression of a BTM’s genes, for each sample in our study. Subacute COVID-19 samples had significantly lower BTM gene scores compared to pre-pandemic controls for several modules related to inflammation, including “Immune Activation”, “TLR and Inflammatory Signaling”, and “Cell Cycle Transcription”; “Enriched in Monocytes” approached significance (Figure 1D, Supplementary Fig. 1). We also used BTM gene scores to explore participant level changes in Hox gene and integrin interaction related modules. Due to overlapping genes across all six Hox gene BTM clusters, we generated a singular BTM gene score for Hox clusters I-VI for each participant sample, which took into account the unique genes across all six modules (Figure 1E). Consistent with the module analysis, gene scores for the “Hox clusters I-VI” and “Integrin Cell Surface Interaction” modules were significantly upregulated in subacute COVID-19 participant samples compared to control samples (Fig 1E and Supplemental Fig. 1D). Overall, participant-level BTM gene scores corroborated our findings from module analysis. Subacute COVID-19 was associated with depleted immune activity and upregulated Hox gene and integrin interaction pathways in SAT.

Next we explored whether the differentially expressed BTMs we identified were associated with specific cell types within human SAT. Consequently, we turned to publicly available single cell RNA sequencing (scRNA-seq) data generated from the stromal vascular cell (SVC) fraction of uninfected human subcutaneous adipose tissue (12). The SVC consists of pre-adipocytes, endothelial, and resident immune cells within adipose tissue but excludes adipocytes. Using the scRNA-seq data, we generated a Uniform Manifold Approximation and Projection (UMAP) which clusters cells by similarity in their overall gene expression and allows us to visualize distinct cell populations (Figure 2A). Mapping all genes included within the “Immune Activation” BTM onto the scRNA UMAP revealed that this transcription module was strongly associated with both macrophages and pre-adipocytes in human SAT (Figure 2B). Thus, the low Immune Activation BTM gene scores suggest that macrophage and pre-adipocyte function or population size may be depleted in subacute COVID-19. S1008A is a gene known to be associated with myeloid cell activation and was also identified as a significantly downregulated DEG in subacute COVID-19 samples (Figure 1B). Indeed, when mapping this myeloid marker onto the scRNA-seq UMAP, its expression appears to be specific to the macrophage subpopulation, suggesting that tissue resident macrophages have decreased inflammatory signatures following COVID-19 infection (Supplementary Fig. 2). Meanwhile, mapping genes related to BTM Hox cluster pathways I-VI showed that expression of these transcription modules is specific to pre-adipocytes, indicating a perturbation to pre-adipocyte function or population size (Fig. 1C).

### 4.4 LC was not associated with significant transcriptomic changes in adipose tissue

Given that prior COVID-19 was associated with changes in the transcriptional landscape of adipose tissue, we next explored whether changes in transcription were associated with LC symptoms. We therefore compared individuals with LC (n=8) to those who were classified as indeterminate by RECOVER score (n=10) (Table 1). PCA of all genes did not distinguish LC from indeterminate samples (Figure 3A). As there was evidence of a batch effect for samples sequenced at different times (Supplemental Figure 3), we used MetaIntegrator to identify DEGs across the two batches (Supplementary Table 1). There were no significant DEGs identified between LC and the indeterminate control samples (39) (Fig. 3B). To specifically investigate the depleted immune activity and Hox cluster BTMs that were associated with subacute COVID-19, we generated gene scores for the “Immune Activation” and “Hox Clusters I-VI” BTMs for all samples in cohort 2. These modules showed no significant difference between LC and the indeterminate sample controls (Figure 3C).

**Figure 3.**
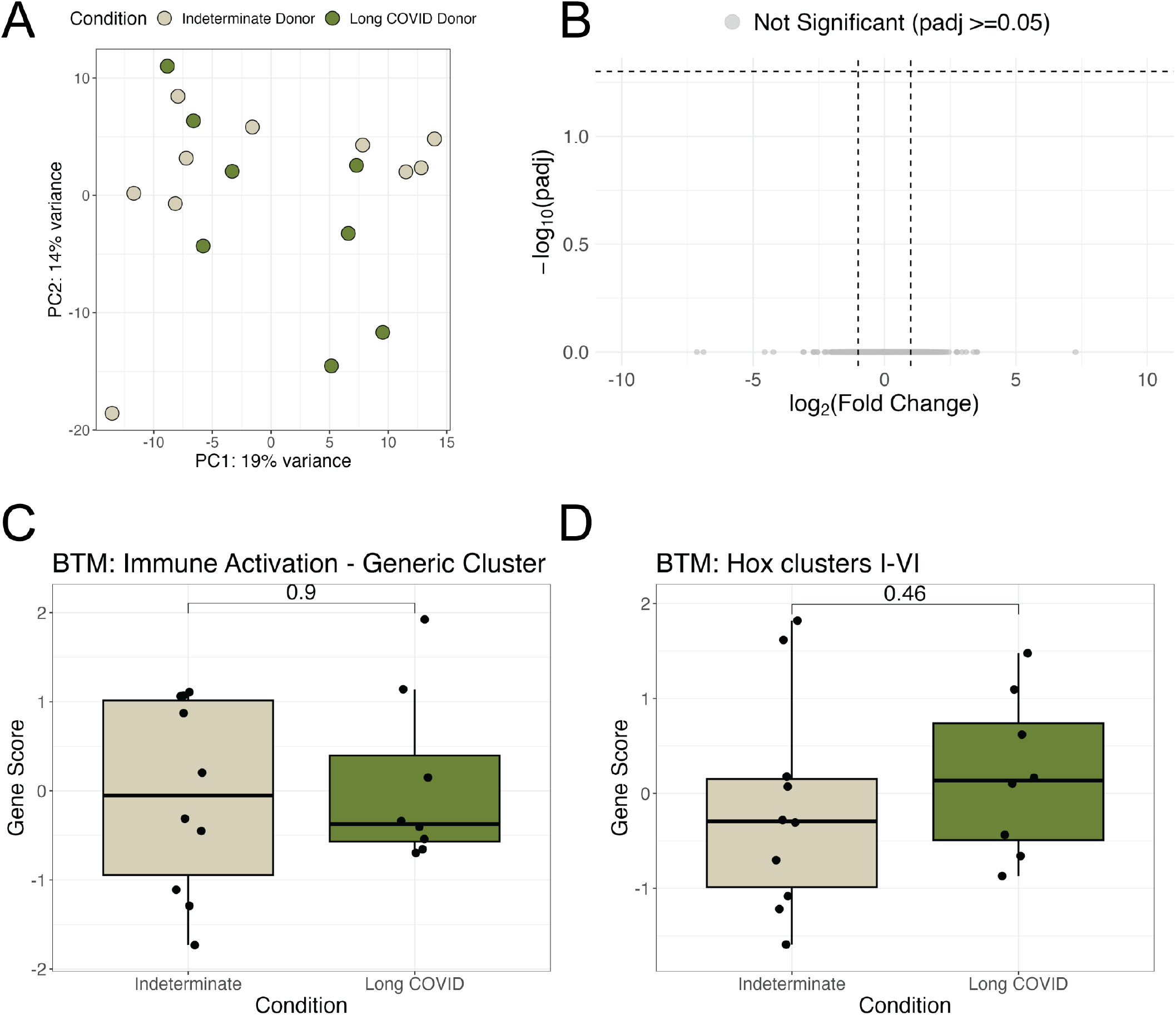
Long COVID is not associated with significant transcriptional changes compared to those with Indeterminate symptoms. **(A)** Dimensionality reduction by principal components analysis based on log2 normalized counts of all genes for participant samples in cohort 2 (Long COVID and indeterminate Status participant samples). Each data point corresponds to the sample of one participant. PC1, principal component 1; PC2, principal component 2; percentage expresses contribution to the overall data variability **(B)** Volcano plot visualization of the differentially expressed genes between Long COVID participants and indeterminate control samples. Horizontal dashed line represents padj = 0.05 and vertical dashed lines represent an absolute log fold change of 1. **(C-D**) Wilcoxon rank-sum p-values comparing gene scores between Control versus Long COVID-19 samples. For boxplot visualizations, the top box line indicates the 75th quartile gene score; middle box line indicates the mean gene score, and the bottom box line represents the 25th quartile gene score.

Finally, we performed immunohistochemistry on a subset of adipose tissue from cohort 2 that had a large difference in severity score (Supplemental Table 2) to determine whether LC affected macrophage density or polarity. We identified inflammatory macrophages as CD68 and CD86 positive cells and anti-inflammatory macrophages as CD68 and CD206 positive cells. We did not detect changes in either type of macrophage, consistent with the lack of transcriptomic changes (Supplementary Fig. 4).

## 5 Discussion

While the mortality of acute COVID-19 has dropped substantially for patient populations with ready access to vaccines, healthcare, and antivirals, LC will continue to place a substantial burden on US and global health systems. Millions of people around the globe are suffering from debilitating symptoms and there remains no cure. Our investigation aimed to explore adipose tissue as a potential reservoir for SARS-CoV-2 persistence and its involvement in systemic dysregulation in subacute and Long COVID phases. Contrary to earlier findings in autopsy studies that identified markers of SARS-CoV-2 persistence in adipose tissue during acute COVID-19 (12,23,40), our results revealed no evidence of persistent viral RNA in SAT samples from either subacute or LC cohorts. These findings challenge the hypothesis that adipose tissue serves as a long-term reservoir for SARS-CoV-2 viral replication.

Though we did not see evidence of virus persistence, prior COVID-19 infection was associated with significant transcriptional remodeling in SAT in the subacute cohort, characterized by downregulation of immune activation pathways and upregulation of homeobox (Hox) genes and integrin-related interactions. Interestingly, the observed depletion of immune-related gene pathways in subacute COVID-19 samples is consistent with earlier studies suggesting that acute SARS-CoV-2 can impair immune cell function and activity in other tissue types, including lung and lymphoid tissues (41,42). Other investigations looking at peripheral blood mononuclear cells (PBMC) found evidence that immune cell dysfunction in acute COVID-19 can be sustained 12-24 weeks post SARS-CoV-2 infection (43). Overall, the transcriptome changes we observed in SAT from subacute COVID-19 participants could point to perturbations in overall adipose tissue function. Considering adipose tissue’s critical endocrine function, we know that even slight changes in SAT function can have systemic effects on systemic inflammation and metabolism in the body (11,15). Due to limited clinical data, we were not able to identify which (if any) subacute COVID-19 participants in our cohort went on to develop LC. As a result we are unable to elucidate whether these changes are evidence of resolving acute COVID-19 or the pathological development of LC. While studies on Hox gene expression in adipose tissue are limited, some investigations involving humans and mice suggest that Hox genes, such as HoxA5, regulate inflammation, energy metabolism, thermogenesis and insulin resistance (44,45). Therefore, inappropriate Hox gene expression may contribute to impaired adipose tissue function (45). Previous investigations have shown integrin activity remains crucial for ECM remodeling, structural stabilization, and repair processes in adipose tissue, especially in the context of obesity-induced inflammation and insulin resistance (46). Indeed, inflammation and fibrosis are characteristic of individuals who exhibit insulin resistance and diabetes (47,48). Therefore, upregulation in integrin activity seen within subacute COVID-19 participants adipose tissue could point to promotion of a pro-inflammatory and fibrotic state, potentially contributing to the development of hyperglycemia and diabetes that are increased following acute COVID-19 infection (15,17,19) as a consequence of adipose tissue changes that promote insulin resistance(49).

Several epidemiological studies have linked COVID-19 infection to an increased risk of developing pro-inflammatory metabolic disorders, namely Type 2 diabetes (T2D), 90 days or more post SARS-CoV-2 infection (15,17,19). Knowing that subcutaneous adipose tissue plays a key role in regulating systemic metabolism and inflammation, we hypothesized that we might find evidence of persistent SAT dysfunction in LC participants and that these perturbations might help distinguish LC participants from those that experience little to no persistent symptoms (indeterminate status participants). Transcriptome analysis of SAT samples did not distinguish participants with significant LC symptoms from indeterminant subjects. These findings suggest that SAT remodeling does not play a major role in driving Long COVID sequelae.

Our study has several limitations. Both cohorts had a small sample size, limiting statistical power and generalizability. Additionally, all participants in cohort 1 had a relatively high BMI, restricting the applicability of these results to lean individuals, and we did not have longitudinal clinical data, so we cannot determine whether these individuals developed LC. In cohort 2, there were 18 individuals who agreed to biopsy from the larger cohort of 60 participants who had been previously infected with SARS-CoV-2, but those 18 were not matched for age or BMI. An additional limitation is that LC symptoms change over time, so individuals may change classification. Finally, the samples were collected and processed at different times, leading to potential batch effects that further complicated the interpretation of findings and precluded direct comparison of the two cohorts. These limitations underscore the need for larger, more controlled studies to validate our observations.

In conclusion, our findings suggest that subacute SARS-CoV-2 infection induces significant transcriptional changes in SAT, particularly in immune-related and integrin pathways. However, there was not a distinct transcriptional signature associated with LC. While adipose tissue does not appear to serve as a reservoir for SARS-CoV-2 replication, its remodeling during subacute COVID-19 may have systemic implications. Future studies should investigate the long-term metabolic and inflammatory consequences of these changes and explore other potential tissue reservoirs or systemic mechanisms underlying Long COVID.

## 5.1 Ethics Statement

Study design and procedures related to recruitment, collection, and assessment of all samples and study participants were approved by institutional review boards. All participants gave written and informed consent.

## 5.3 Author Contributions

CAB, TM, CR, JZL, PAK, BSF conceived the work and obtained funding; SD-C performed experimental studies and coordinated the study; HC, KR, UMM, TRB, HM, BSF, ZL, HM, BML, GEE, and PAK performed experimental studies; SJS, NT, JY, NS, KMA, EMA, IE, HM, AC enrolled participants and performed clinical assessment; TML, SC performed biopsies; SD-C, KR prepared the figures; SD-C, KR, CAB wrote the manuscript. All authors contributed to the manuscript and approved the submitted version.

## 5.5 Acknowledgments

We would like to thank all research participants for their contribution to our research and Researching COVID to Enhance Recovery (RECOVER), a National Institutes of Health (NIH) research initiative for funding our work. We would also like to thank Drs. Arjun Rustagi and Giovanny Martínez-Colón for input, protocols, and advice.

## 5.6 Data Availability Statement

Raw sequencing data will be deposited under the NCBI Bioproject [Insert ID] and processed data under GSEXXX upon publication. Analysis scripts will be available at https://github.com/BlishLab/2024_COVID19_Adipose upon publication.

## 6 Scope Statement

Our manuscript explores the role of adipose tissue as a potential reservoir for SARS-CoV-2 and its involvement in Long COVID pathophysiology. Long COVID, characterized by symptoms persisting beyond 90 days post-infection, remains poorly understood. To investigate, we analyzed subcutaneous adipose tissue (SAT) from two cohorts: participants in the subacute COVID-19 phase (28–90 days post-infection) and those with Long COVID (>90 days post-infection). No evidence of persistent SARS-CoV-2 RNA was detected in SAT from either cohort. However, significant transcriptional remodeling in the subacute COVID-19 group, compared to pre-pandemic controls, suggested immune cell exhaustion and altered tissue function. Conversely, no consistent gene expression changes were observed in SAT from Long COVID participants compared to Indeterminate status participants, indicating a limited role for adipose tissue in sustaining Long COVID symptoms. These findings highlight disruptions to SAT immune activity and tissue function following SARS-CoV-2 infection, providing insights into inflammatory dysregulation during recovery. This study aligns with *Frontiers in Endocrinology* by advancing understanding of metabolic and immunological factors in viral diseases and their implications for long-term health, addressing key themes within the journal’s focus on endocrine dysfunction and innovative therapies.

