## Supplementary Figures for "COVID-19 induces persistent transcriptional changes in adipose tissue that are not associated with Long COVID"

### *Supplementary Material*

**Supplementary Table 1:** Clinical Data for All Donors in Cohort 1 (Subacute COVID & Pre-Pandemic Controls Samples) and Cohort 2 (Long COVID & indeterminate Status Controls. *Ct Value 37 or higher is considered below the limit of detection; Undetermined is considered a Ct value greater than 40. **0= no symptoms to 20= Most Severe.

**Supplementary Figure 1:**

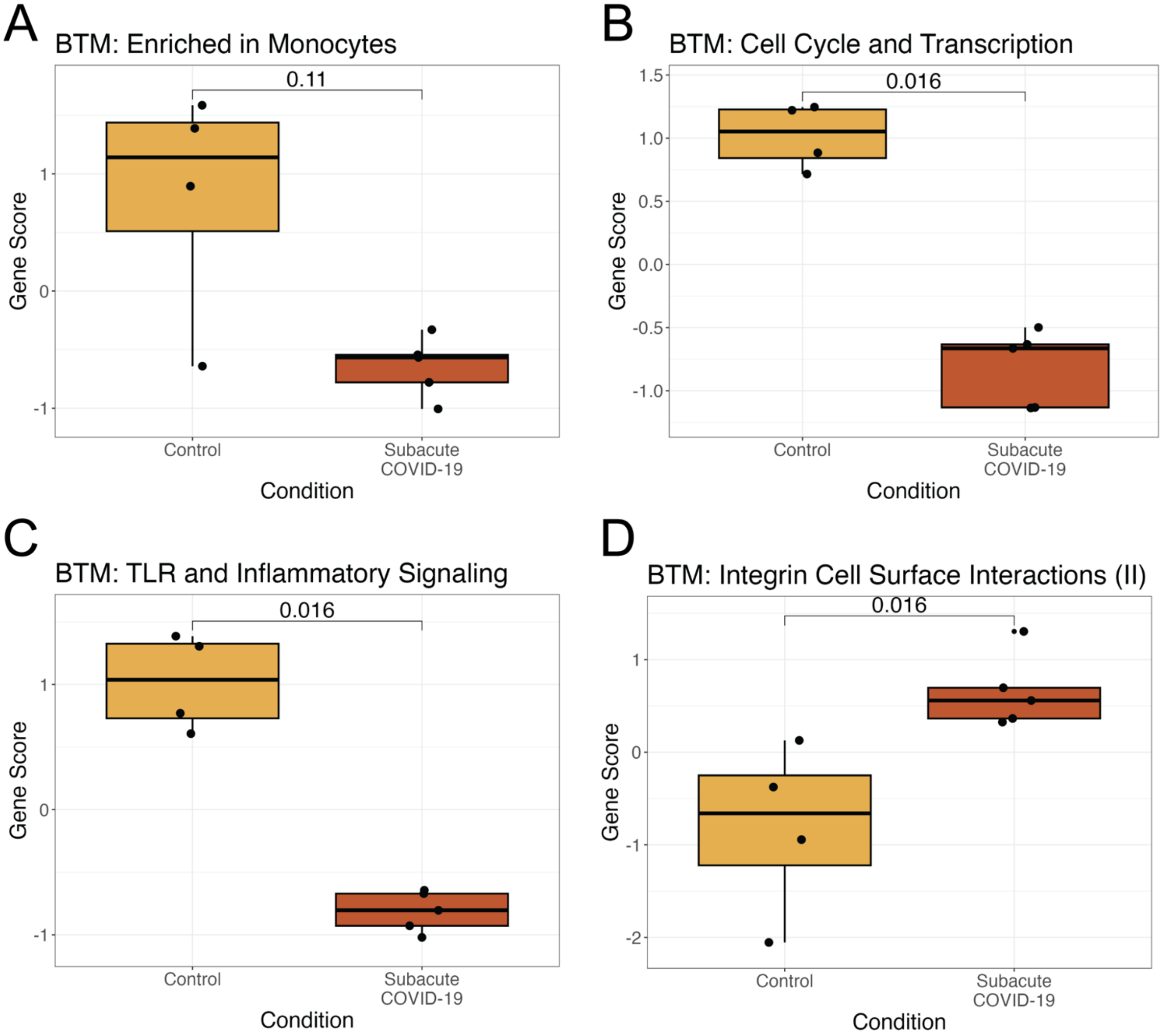


**Supplemental Figure 1 Additional BTM Gene Scores for Cohort 1**. Box plots of "Enriched in monocytes" **(A),** "Cell cycle and transcription" **(B)**, "TLR and inflammatory signaling" **(C)**, and "Integrin Cell Surface Interactions (II)" **(D)** BTM gene scores for subacute COVID-19 samples (n=5) and pre-pandemic control samples. The top box line indicates the 75th quartile gene score; middle box line indicates the mean gene score, and the bottom box line represents the 25th quartile gene score. All p-values generated with Wilcoxon rank sum test.

**Supplemental Figure 2**

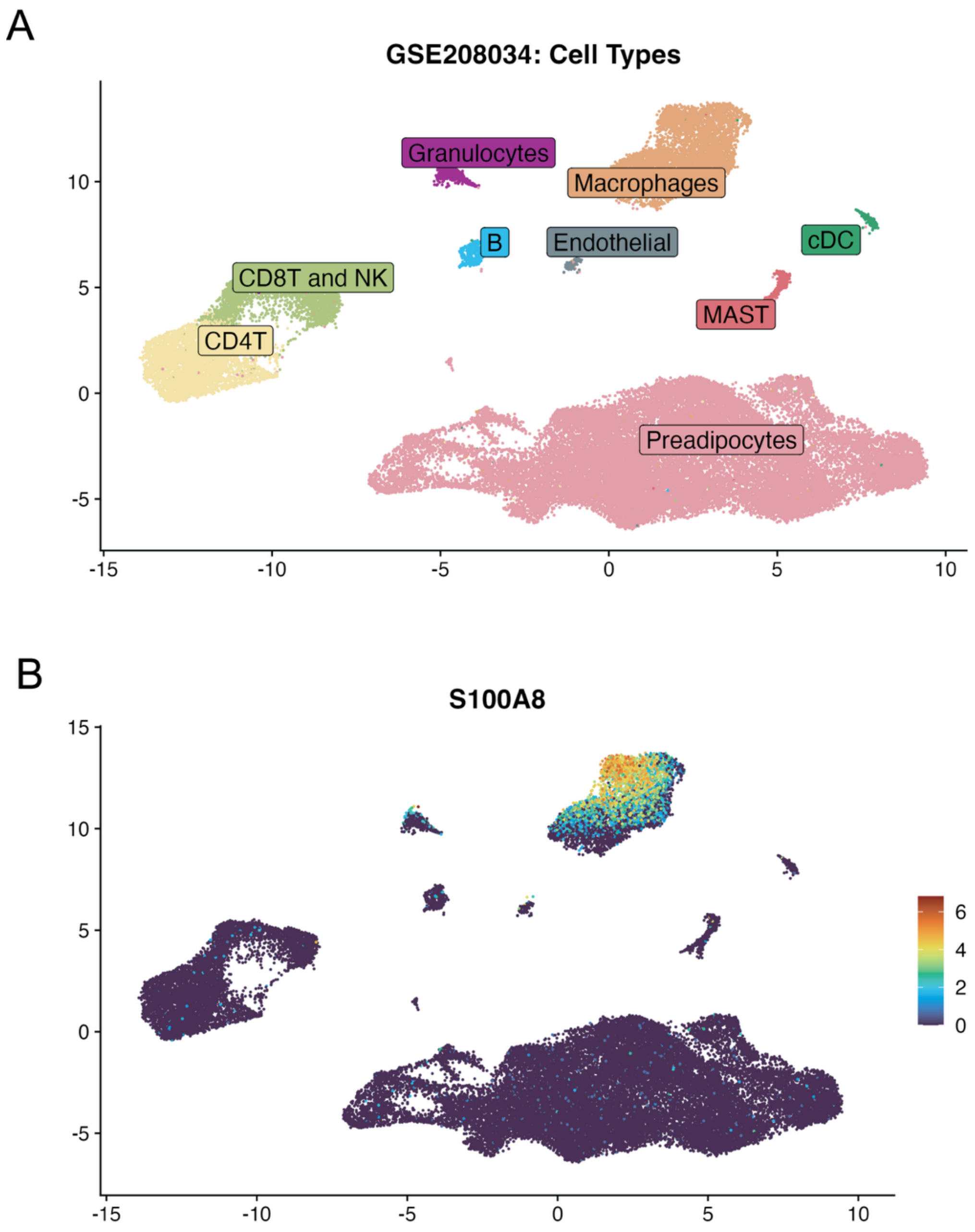


**Supplemental Figure 2. Expression of S100A8 is Strongly Associated with Resident Tissue Macrophages in SAT (A)** Uniform Manifold Approximation and Projection (UMAP) of Single Cell RNA (scRNA) dataset from Stromal Vascular Fraction (SVF) of human SAT (GSE208034) colored by cell type (annotated using Seurat), repeated from Fig. 2A for easier reference **(B)** UMAP of the expression of **S100A8** across cell types in SVF of human SAT (GSE208034) colored by cell type (annotated using Seuratv5).

**Supplemental Figure 3**

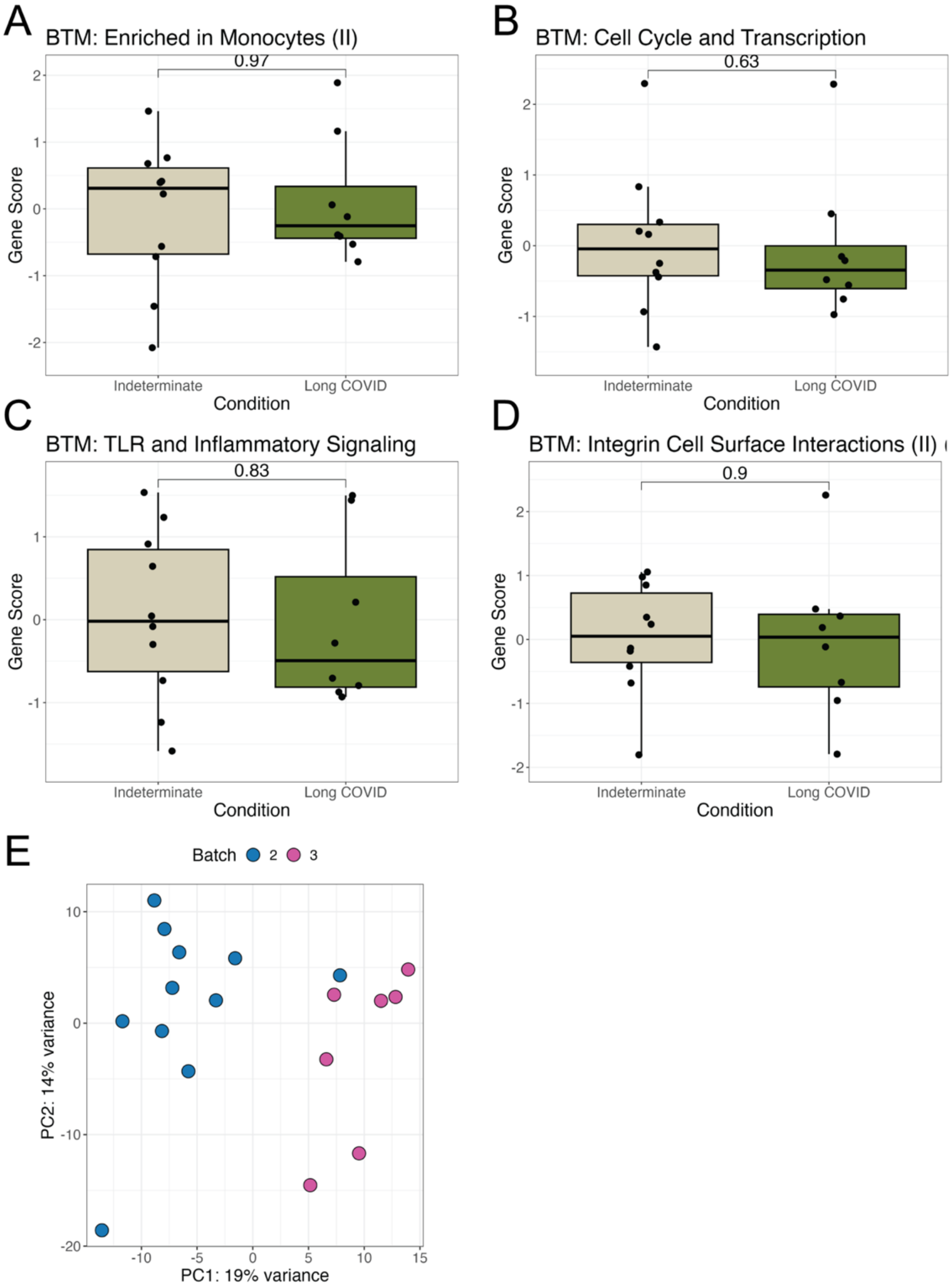


**3**

**Supplemental Figure 3. Additional BTM Gene Scores and PCA for Cohort 2**. Box plots of "Enriched in monocytes" **(A),** "Cell cycle and transcription" **(B)**, "TLR and inflammatory signaling" **(C),** "Integrin Cell Surface Interactions (II)" **(D)** BTM gene scores for Long COVID-19 samples (n=8) and indeterminate status samples (n=10). The top box line indicates the 75th quartile gene score; middle box line indicates the mean gene score, and the bottom box line represents the 25th quartile gene score. All p-values generated with Wilcoxon rank sum test. **(E)** Dimensionality reduction by principal components analysis based on log2 normalized counts of all genes for participant samples in cohort 2 (LC and indeterminate Status participant samples). Each data point corresponds to the sample of one participant and is color coded by the RNA sequencing batch number. PC1, principal component 1; PC2, principal component 2; percentage expresses contribution to the overall data variability

**Supplemental Figure 4**

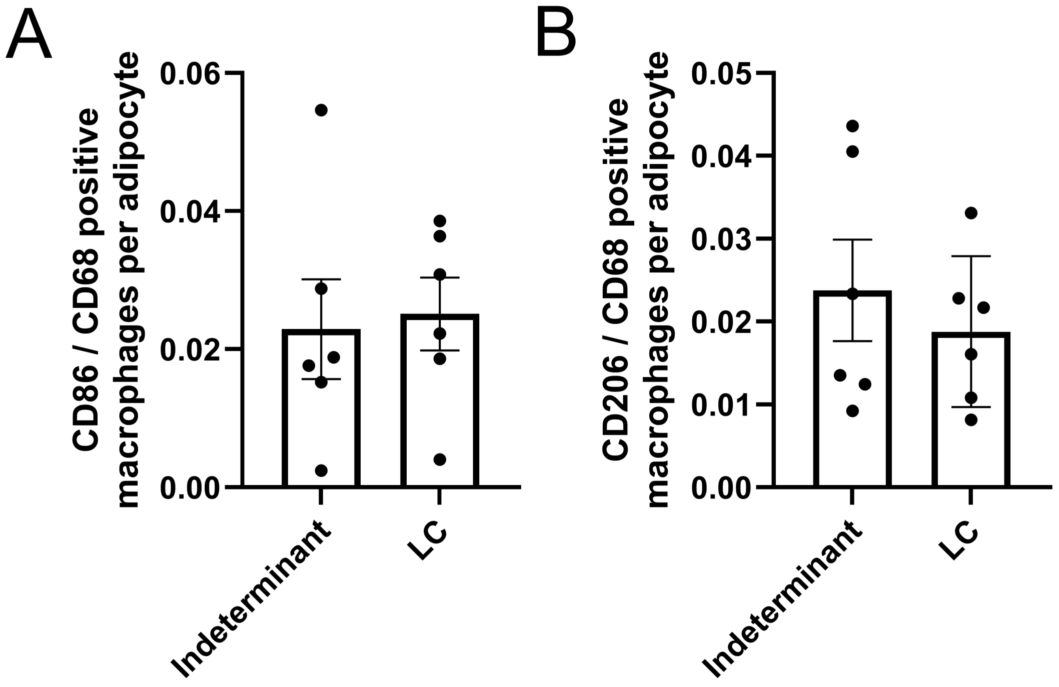


**Supplemental Figure 4. Long COVID-19 does not Affect SAT Macrophage Density or Polarity**. Subcutaneous adipose was co-stained with CD86 and CD68 (A) or CD206 and 68 (B). Data represent means ± SEM (Indeterminant (-) n=4; LC (+) n=8) and were analyzed by an unpaired, 2-tailed Student's t-test.

**Supplementary Table 2**

| | Indeterminant | Long Covid |
|---|---|---|
| Number of Samples (Male/Female) | 6 (1/5) | 6 (1/5) |
| Mean Age | 53.7 ± 5.2 | 55.5 ± 7.1 |
| Mean BMI | 27.5 ± 2.3 | 32.1± 1.9 |
| Mean Recover Severity Score | 2.3 ± 1.1 | 12.8 ± 1.3 |

**Supplementary Table 2:** Baseline characteristics of research participants in which SAT macrophages were quantified by immunohistochemistry.
